# Delivering biochemicals with precision using bioelectronic devices enhanced with feedback control

**DOI:** 10.1101/2023.08.29.555386

**Authors:** Giovanny Marquez, Harika Dechiraju, Prabhat Baniya, Houpu Li, Maryam Tebyani, Pattawong Pansodtee, Mohammad Jafari, Mircea Teodorescu, Marco Rolandi, Marcella Gomez

## Abstract

Precision medicine tailors treatment in a way that accounts for variations in patient response. Treatment strategies can be determined based on factors such as genetic mutations, age, and diet. Another way of implementing precision medicine in a dynamic fashion is through bioelectronics equipped with real-time sensing and intelligent actuation. Bioelectronic devices such as ion pumps can be utilized to deliver therapeutic drugs. To be able to perform precision medicine, medical devices need to be able to deliver drugs with high precision. For this, closed-loop control is required to be able to change the treatment strategy as new information about the response and progression of the biological system is received. To this end, a sliding mode controller is utilized given its ability to perform satisfactory control actions when there is model uncertainty. The controller is used in an experiment with the goal of delivering a pre-determined dosage of fluoxetine throughout a period of time.

## Introduction

Recent developments in bioelectronics hold promise for advancing precision medicine [1]. Bioelectronic devices have been created that allow for monitoring blood sugar levels, controlling stem cell fate, applying electrical stimulation, and delivering therapeutic drugs [2–4]. Ion pumps are bioelectronic devices that allow for the movement of ions and charged drugs to move toward a targeted area electrophoretically [5]. They have applications in precision medicine when applying a concentration of a drug at a specific region and time is more effective than passive drug delivery methods such as digested pills [6].

Such applications require the ability to carefully control the delivery of therapeutics with precision. In [7], the authors discuss the ability of feedback control to enhance the capabilities of bioelectronic devices. However, implementing feedback control in bioelectronic devices presents unique challenges including variability in device performance. It is difficult to fabricate bioelectronics in a way that the end process is exactly the same. Due to variations occurring in the fabrication process, the bioelectronic devices can differ in their properties. Components in the device can also vary in their performance when the device is used for an extended period of time [8]. This can lead to differing responses across devices and time when used in an experimental setting.

To address the above-mentioned challenges, in previous work, a neural network (NN)-based machine learning algorithm was used with ion pumps to successfully control the pH level of a target solution [9]. A feedback control algorithm was able to regulate pH levels according to a target time-varying trajectory. In [10], the authors were able to use a similar control algorithm to control the membrane potential of stem cells by way of regulating pH of the extracellular environment. Thus, an adaptive feedback control algorithm helps to mitigate variations in system response even when they evolve in time.

The second primary challenge in the control of bioelectronic devices is their limited operating range. That is, the voltages that can be applied to the device are limited.

This causes a saturation issue that needs to be taken into account. In [11], the authors presented an alternative approach to arriving at an adaptive algorithm that has the benefits of the NN-based controller but explicitly handles saturation with guaranteed convergence. This algorithm was tested *in silico*.

In this paper, we present an adaptation of the algorithm presented in [11] to control the amount of biochemical being delivered by an ion pump. In our adaptation, we apply a heuristic switching algorithm for modification of the controller gains to improve performance. The biochemical chosen is fluoxetine, which has been shown to be a relevant drug in wound healing in diabetic mice [12]. We demonstrate the algorithm with three different reference signals that result in the delivery of the same total drug concentration over the period of actuation. This was done to show the controller’s ability to keep the current at the reference regardless of the signal used as well as to allow for different methods of delivery. We demonstrate that the controller is able to perform at a reasonable level, keeping the current near the reference throughout the experiment.

The paper will be structured as follows. Subsection “Bioelectronic Ion Pump” will introduce the ion pump and its capability to deliver flouxetine. Subsection “Controller” will discuss the sliding mode controller being used. Subsection “Experimental Set up” will discuss the experimental setup and how things are integrated together. Sections “Results” and “Discussion” will show the results and a discussion on them.

## Materials and methods

In this section, we introduce the ion pump and discuss its ability to deliver fluoxetine. We then discuss the sliding mode controller and give a brief outline of how the controller works. Finally, we present an overview of the experimental setup used.

### Bioelectronic Ion Pump

Ion pumps are bioelectronic devices that can perform precise electrophoretic delivery of ions and molecules from a region of high concentration in the device, known as the reservoir, to the region of interest, known as the target [5, 13]. Over the years, ion pumps have been used to modulate pH in a local environment [14], treat neurological disorders [15, 16], induce macrophage recruitment [17] and even to deliver biochemicals in plants [18]. In this work, we use the ion pump to deliver Fluoxetine, a biochemical which is a selective serotonin reuptake inhibitor known to have immunomodulatory effects [12].

The ion pump transfers the biochemical from the reservoir to the target, with a voltage (V_pump_, typically between 0.5*V* and 2*V*) applied between the working electrode (Ag) and the reference electrode(Ag/AgCl) (see Fig 1(a)). A negatively charged hydrogel selectively transports the biochemical between the target and the reservoir. In the reservoir, we have 0.01*M* of fluoxetine hydrochloride. Fluoxetine exists as a positively charged species at a pH of 5. For a positive V_pump_, the Fluoxetine moves through the hydrogel-filled capillary and is transported to the target. Fig 1(b) shows an image of the device in a six-well plate with a buffer solution. This acts as our target for the experiments.

**Fig 1.**
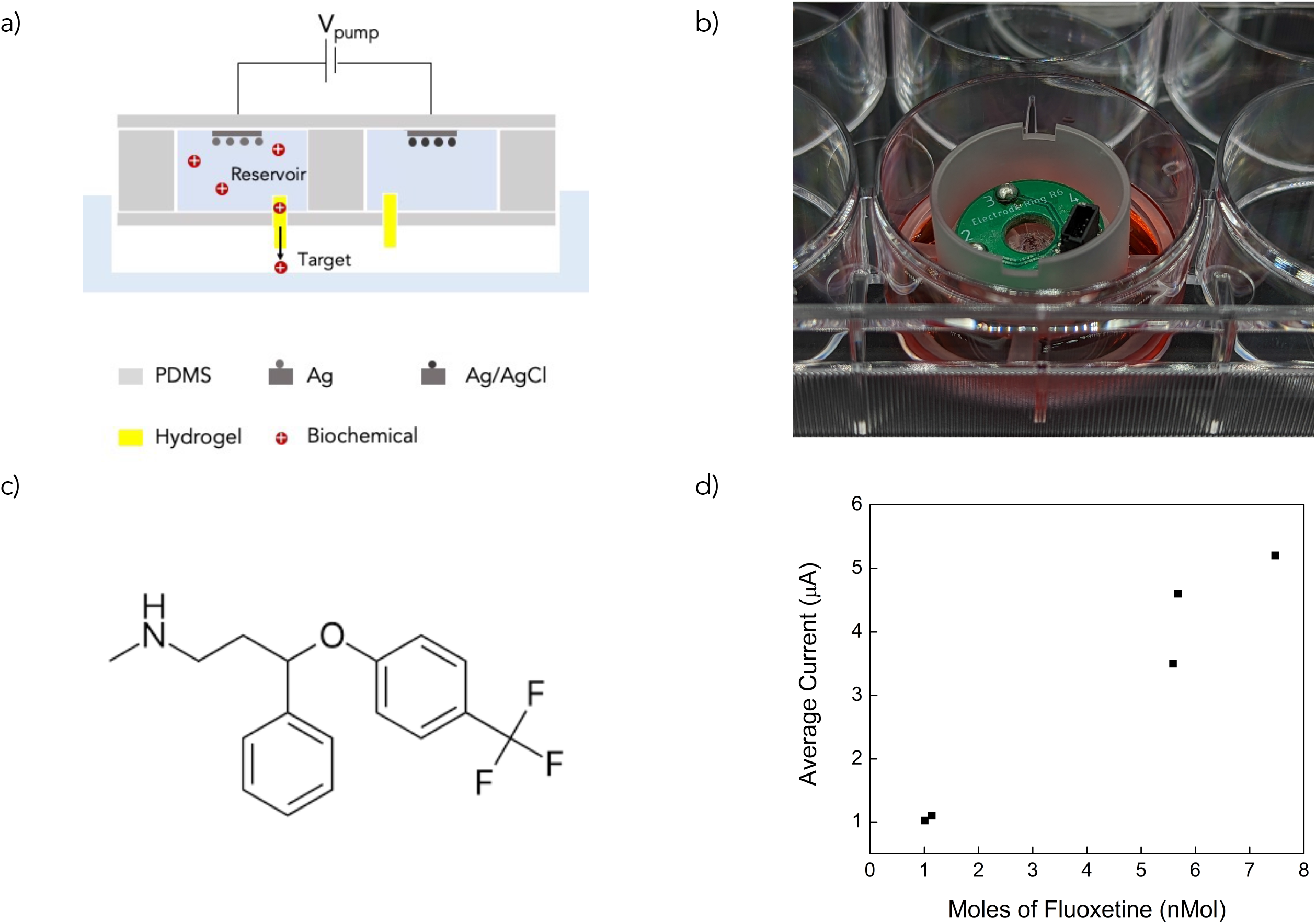
(a) Schematic of the bioelectronic ion pump. (b) Image of the ion pump in a 6-well plate with buffer solution. (c) Chemical structure of Fluoxetine (d) Graph showing the relationship between the current produced by the ion pump and the amount of fluoxetine delivered.

We collect the target solution after completion of delivery and use High-Performance Liquid Chromatography (HPLC) to estimate the amount of Fluoxetine in the target. We establish a relationship between the current produced during the biochemical delivery and the corresponding amount of fluoxetine delivered to calibrate the ion pump’s performance (see Fig 1(d)) Based on these results, we found the devices to be delivering the biochemical with an efficiency of approximately 20%, where efficiency is defined as follows

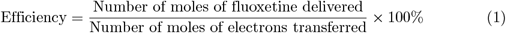

### Controller

Sliding mode control is known for its ability to handle system uncertainties including unmodelled dynamics [19]. A sliding mode controller attempts to achieve a reference signal by forcing the system to slide toward a stable manifold. The controller drives the system towards a designed sliding manifold, where the system stays, sliding along the manifold until it reaches equilibrium. One major drawback with the control design is the chattering of the control signal that tends to appear [20]. However, researchers have developed many methods to address this so the control method can be used in real-world applications. It is applied in robotics, process control, induction motors, and power converters [21–23].

A similar sliding mode controller developed in [11] is utilized for the output feedback control experiment in this paper (see Fig 2). The controller developed in [11] was shown to perform well with a mechanistic model of an ion pump that included saturation. The challenges encountered in simulations are comparable to the ones expected in these experiments. This inspired confidence that the controller developed in [11] would be a viable controller.

**Fig 2.**
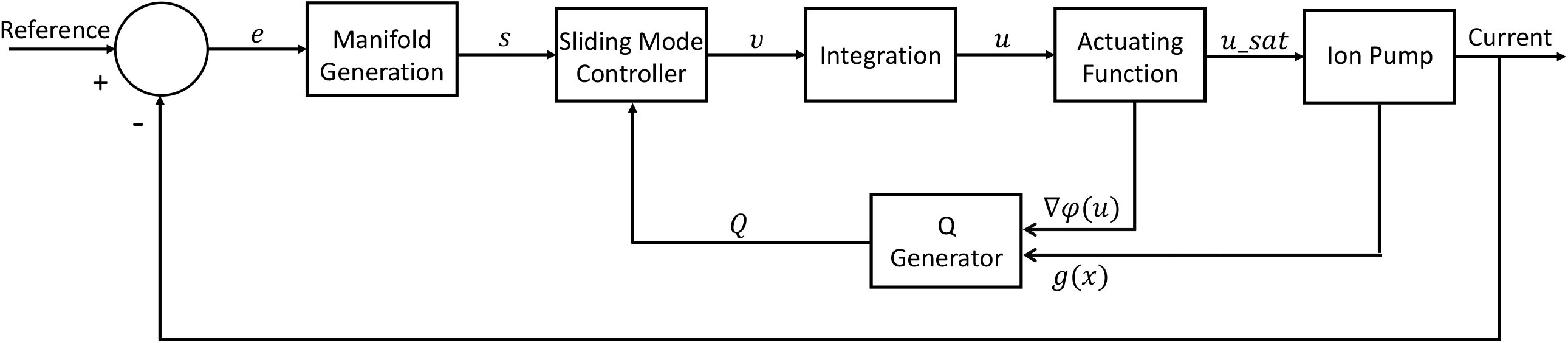
Closed-loop control architecture for automated ion pump actuation to achieve the desired current value in order to control the concentration of fluoxetine.

To begin, we consider our system to be an affine nonlinear system with input saturation represented by the following state space model

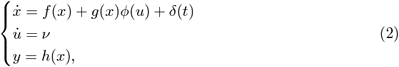

where *x ∈* ℝ^*n*^ is the state vector. In our experiment, *x* is the current being read from the system. An artificial control signal *ν∈ ℝ*^*m*^ is introduced to help reduce the known chattering issue with sliding mode controllers. The output of the function sat(*ϕ*(*u*)) *∈* ℝ^*m*^ is the control input vector. In this application, this is the voltage applied to the system. The variable *y∈* ℝ^*n*^ is the output vector of the system. In the experiment here *y* = *x* which is the read current from the system. The function *f* (*x*) *∈* ℝ^*n*^ is an unknown locally Lipschitz nonlinear function and *g*(*x*) *∈* R^*n×m*^ is an unknown input coefficient value. The time-varying variable *δ*(*t*) *∈* ℝ is a sufficiently smooth disturbance occurring throughout the experiment. We design the manifold as

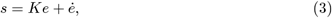

where *K* is a positive constant, and *e* is the error defined as

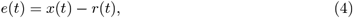

*x*(*t*) is the current value, and *r*(*t*) is the desired reference value for the current.

The sliding mode controller then calculates the artificial control input to be used for calculating the actual control signal to be sent to the ion pump

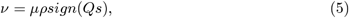

where *ρ* is a positive coefficient and *Q* is obtained as

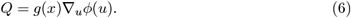

We require the sign of *g*(*x*) found through experiments to be *μ* = *−*1, which is the case when the response of the system is monotonically increasing with the input. In some cases, the rate of change in the response could be faster or slower in one direction than the other. Therefore, we introduce a heuristic switching algorithm 1 for better adaptation to these conditions.

#### Algorithm 1 Gain Update Algorithm

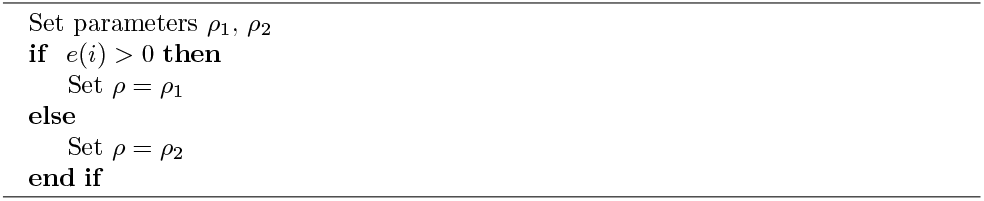

The actuating function used here is defined as

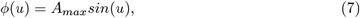

where *A*_*max*_ is the max/min voltage allowed by the ion pump. For the experiments, we set *A*_*max*_ = 3. To help reduce the chattering signal that tends to occur with sliding mode controllers, *ν* is integrated to get *u*. Integration is done using the trapezoidal rule where we have a sampling time of *T* = 1. It should be noted that using the trapezoidal rule causes us to start the algorithm at *t* = 2. Finally, to make sure we stay in the saturated region, we applied *ϕ*(*u*) to get the final voltage that will be applied to the ion pump

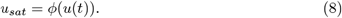

Algorithm 2 shows a detailed description of the implementation of the developed algorithm.

For details regarding the stability proof of the developed controller, readers are referred to our previous work [11].

#### Algorithm 2 Sliding Mode Control Algorithm

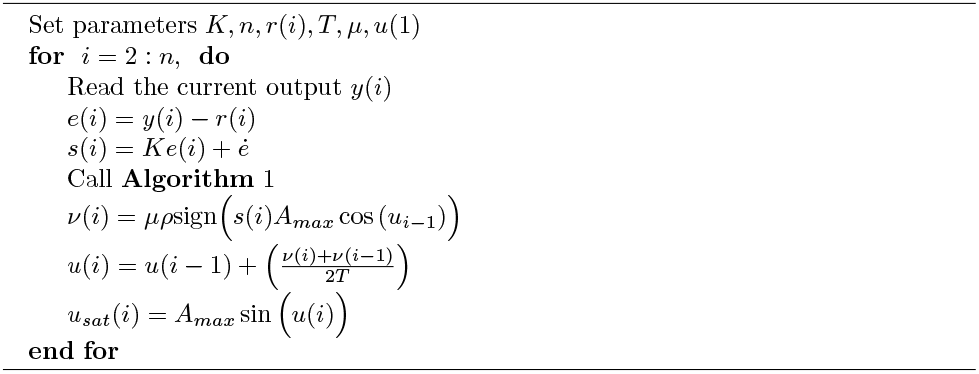

### Experimental Set up

The ion pump is integrated with a printed circuit board (PCB) interface which connects the electrodes in the pump to an external power supply. We use silver pins and a conductive silver paste to connect the Ag/AgCl electrodes in the reservoir to the PCB which is then soldered to the ion pump. Jumper cables are used to connect this PCB to a Raspberry Pi-based external voltage controller which can connect to the WiFi [24]. The device is seated in a six-well plate with a buffer solution acting as the target for delivery. The external voltage controller applies the control commands it receives from the sliding mode controller to the ion pump. The setup is able to successfully apply up to 16 different voltages with a resolution of 1.95*mV* per its channels. In addition, it is able to successfully read up to 16 current values from the ion pump but at a potential sacrifice in the accuracy of the reading, where the resolution of the current reading is 0.125*nA*. This could lead to extra uncertainty in the system.

Fig 3 shows the closed-loop experimental setup. The error is calculated using the value of the current output read from the ion pump and the desired reference value. The sliding mode controller evaluates the next voltage value to drive the current toward the desired reference. This is sent to the Raspberry Pi through a WiFi connection between the Raspberry Pi and a laptop where the control algorithm is running. The Raspberry Pi applies the voltage value to the ion pump through a connected cable. This closes the loop.

**Fig 3.**
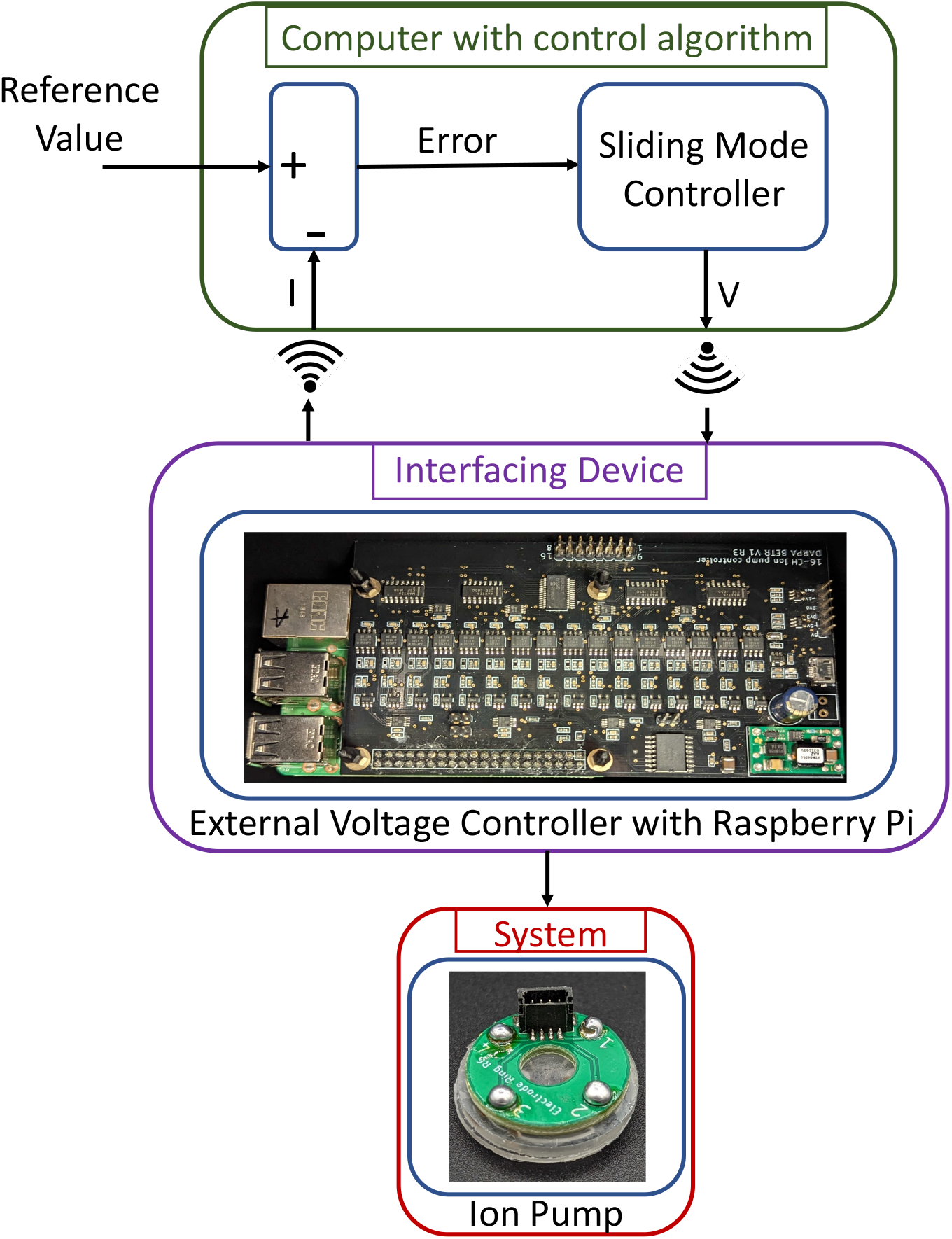
Schematic of the experimental closed-loop setup. The error is computed using the values of the current output read from the ion pump and the desired reference value. The sliding mode controller evaluates the next voltage value. This is sent to the external voltage controller with Raspberry Pi through a WiFi connection with the laptop running the control algorithm. The external voltage controller applies the value through a connected cable from the ion pump. After the voltage has been applied, the external voltage controller reads the current from the ion pump and sends it back to the laptop with the control algorithm through WiFi closing the loop.

## Results

Fig 4 shows experimental results for feedback control on current in the ion pump device using sliding mode control. In the first column, the blue lines indicate the desired current set as a reference for the feedback control algorithm. The red line is the current being measured from the device in real-time. The second column shows the controller output (i.e., voltage) being applied to the ion pump in green. The third column shows the corresponding tracking error in cyan.

**Fig 4.**
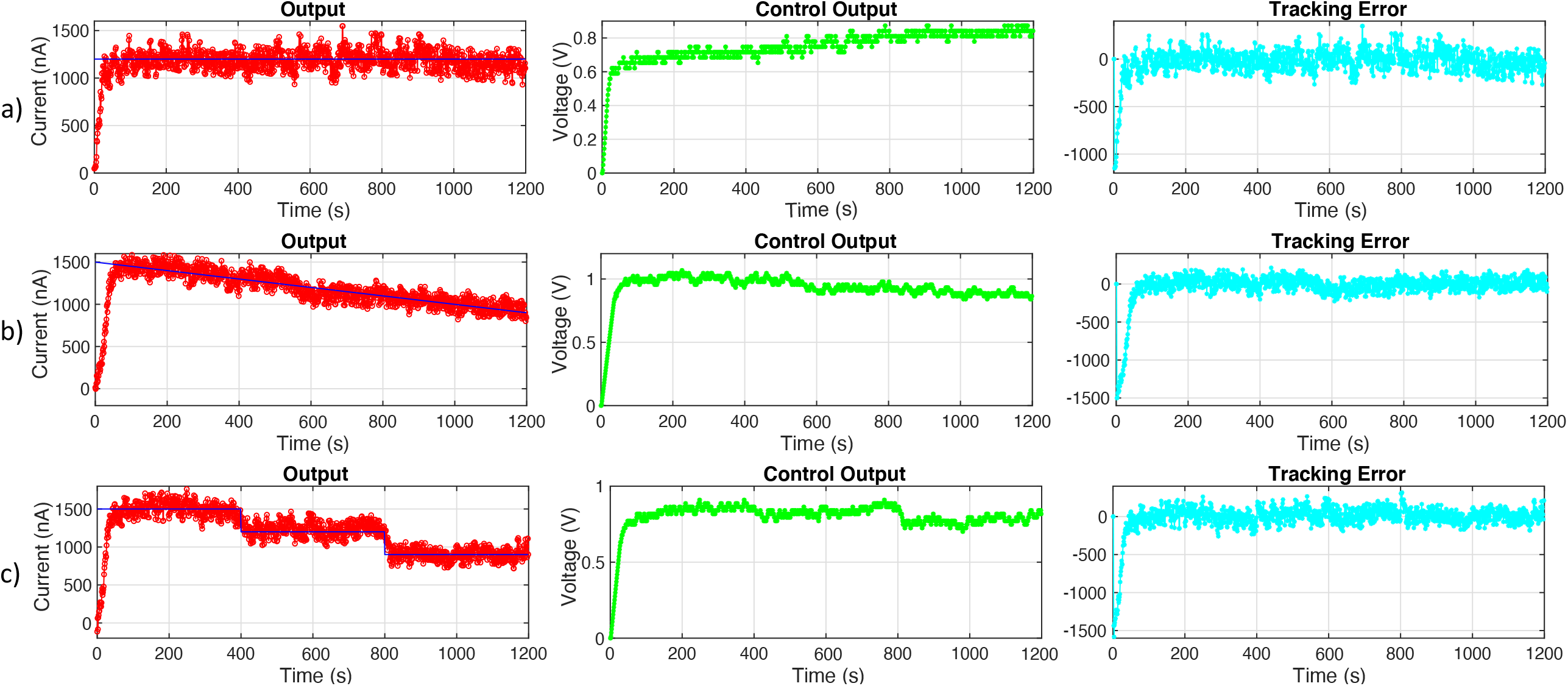
Experimental results for feedback control on current in the ion pump device using sliding mode control. Row (a) shows the ion pump response to a constant reference signal at 1200*nA*. Row (c) shows the ion pump response to step changes in the reference signal starting at 1500*nA* dropping by 300*nA* each 400 seconds. Row (b) shows the ion pump response to a gradual decline reference signal beginning at 1500*nA* and ending at 900*nA*.The blue line in the first column indicates the desired current set as a reference for the feedback control algorithm. The red curve represents the measured current from the device in real time. The second column shows the control output that is delivered to the device in green. The third column shows the error between the desired and measured current referred to as tracking error in cyan.

We establish a value for the reference signal by estimating the amount of current required to deliver a dosage of 0.45*mg* fluoxetine. Based on device calibration, we estimate that a current of 1.2*μA* (1200*nA*) is required to meet the desired dosage of biochemical.

In the first experimental setup (see Fig 4(a)), we test the controller’s performance on a constant reference signal of 1200*nA*. In the second experiment (see Fig 4(b)), we test the controller’s performance on a gradual descending reference signal starting at 1500*nA* dropping to 900*nA* by the end of the experiment. In the third experiment (see Fig 4(c)), we test the controller’s performance on a step-wise descending reference signal starting at 1500*nA*, dropping by 300*nA* every 400 seconds. Note that the fluctuations around the reference signals at steady-state have a relative error of less than 7% in all the experiments. We define the system to be at steady-state after the first 100 seconds of the experiment to allow for all three runs to reach the reference from the starting current of 0*nA*. Since the amount of biochemical delivered is determined by integrating over the measured current, we don’t expect that the high-frequency fluctuations in current measurements have a high impact on the overall uncertainty in the biochemical delivery. Although the fluctuations make it difficult to keep the current exactly at the reference, the average current throughout each experiment can be calculated to show the desired amount of fluoxetine is being delivered. The averages for each reference signal are as follows: constant - 1171.49*nA*, gradual decline - 1166.45*nA*, decreasing steps - 1184.75*nA*. Table 1 shows the relative error information along with the averages.

**Table 1.** Quantitative measures of all the experiments.

| Experiment | average output signal | average steady-state relative error |
| --- | --- | --- |
| constant | 1171.49nA | 7% |
| gradual decline | 1166.45nA | 6% |
| decreasing steps | 1184.75nA | 7% |

## Discussion

We see that the controller had to gradually increase the voltage applied to the ion pump in order for it to maintain the constant reference. This is due to the device’s decaying performance as the experiment continues. This highlights the need for an adaptive control strategy and reinforces why open-loop control is unable to deliver a desired concentration with precision. One thing to notice is that the average current throughout the experiment was always lower than the actual reference. It’s unclear why this is happening but it can be compensated for by increasing the reference signal slightly past the wanted reference to offset the consistent error being seen.

An enhancement that can be applied to achieve better control of the ion pump is to develop an algorithm that automatically tunes the control parameters. The reason for this is that the efficiency of the ion pump tends to decline over time. This would mean that the response of the device will change over the course of the experiment, which could affect the performance of the sliding mode controller. Adaptive parameters could lead to better performance since they would adjust as needed to the varying response of the ion pump.

Some of the oscillatory behavior of the current could be due to the interfacing board used. When the controller sends a voltage value that is smaller than the resolution of 1.95*mV* then the board is unable to send the exact voltage value. This can cause slight oscillatory behavior with the current as small changes in voltage are able to cause large changes in the current as seen in Fig 4. Although exact control of the current at the reference is not achievable, we see in Table 1 that the average of the current is near the desired value and is within acceptable relative error bounds.

Finally, the presented work combining closed-loop control and delivery of fluoxetine by ion pumps can have real world applications in the wound-healing field. People can respond differently to the same applied medical treatment for reasons such as age, ethnicity, and genetics [25]. This makes it difficult for physicians to treat people optimally. Through precision medicine, custom treatment options can be created depending on an individual’s needs. Closed-loop control allows for the customization of the treatment strategy by changing the strategy in real-time as new information is gained [26]. This can be done by interfacing the feedback control regulated ion pump presented here with a higher-level control algorithm that provides the reference signal [11]. In this way, wound healing might be treated by sensing how well a wound is responding to a specific treatment and changing the dosage of biochemicals as needed in real-time.

## Conclusion

We demonstrated a method for delivering a desired amount of biochemical using closed-loop control. By finding the efficiency of the ion pump in delivering fluoxetine, one can compute the amount of fluoxetine being delivered using the current of the device. A sliding mode controller was used *in vitro* to successfully keep the current at the desired reference which allowed the desired amount of fluoxetine to be delivered.

## Acknowledgments

The authors thank all active collaborators who have engaged us in conversations on the topic of drug delivery and guidance on the experimental setup. The authors would like to thank Bashir Hosseini Jafari with developing the sliding mode controller used.

